# *DCT4* - a new member of the dicarboxylate transporter family in C_4_ grasses

**DOI:** 10.1101/762724

**Authors:** Sarit Weissmann, Pu Huang, Koki Furuyama, Madeline A. Wiechert, Mitsutaka Taniguchi, James C. Schnable, Thomas P. Brutnell, Todd C. Mockler

## Abstract

Malate transport shuttles atmospheric carbon into the Calvin-Benson cycle during NADP-ME C_4_ photosynthesis. Previous characterizations of several plant dicarboxylate transporters (DCT) showed that they efficiently exchange malate across membranes. Here we identify and characterize a previously unknown member of the *DCT* family, *DCT4*, in *Sorghum bicolor*. We show that SbDCT4 exchanges malate across membranes and its expression pattern is consistent with a role in malate transport during C_4_ photosynthesis. *SbDCT4* is not syntenic to the characterized photosynthetic gene *ZmDCT2*, and an ortholog is not detectable in the maize reference genome. We found that the expression patterns of *DCT* family genes in the leaves of *Z. mays*, and *S. bicolor* varied by cell type. Our results suggest that sub-functionalization of members of the *DCT* family for the transport of malate into the bundle sheath (BS) plastids occurred during the process of independent recurrent evolution of C_4_ photosynthesis in grasses of the PACMAD clade. This study confirms the value of using both syntenic information and gene expression profiles to assign orthology in evolutionarily related genomes.

## INTRODUCTION

Three subtypes of C_4_ photosynthesis are generally recognized as defined by the primary decarboxylase in the bundle sheath (BS) cells: chloroplastic NADP-dependent malic enzyme (NADP-ME); mitochondrial NAD-dependent malic enzyme (NAD-ME); and cytosolic phosphoenolpyruvate carboxykinase (PEPCK) (Hatch and Slack, 1966; Hatch, 1971; Rathnam and Edwards, 1977). Different plant species may contain various combinations of these three subtypes (Hatch, 1971; Chapman and Hatch, 1979; Furbank, 2011; Pick *et al*., 2011; Wang *et al*., 2014b). The movement and exchange of malate across membranes, by dicarboxylate transporters (DCTs/DiTs), plays a significant role during photosynthesis in NADP-ME and NAD-ME C_4_ species (Ding *et al*., 2015). In C_3_ plants, DCTs are crucial to nitrate assimilation, such as the GS/GOGAT cycle and photorespiration (Linka and Weber, 2010; Kinoshita *et al*., 2011). Taniguchi *et al.* characterized several plant DCTs that efficiently exchange malate across membranes (Taniguchi *et al*., 2002; Taniguchi *et al*., 2004). The differential expression of C_4_ photosynthesis genes in mesophyll (M) and BS cells (John *et al*., 2014; Tausta *et al*., 2014; Wang *et al*., 2014a) suggests that different malate transporters may be needed to move malate out of the chloroplasts of M cells and into the chloroplasts of BS cells. In *Zea mays*, an NADP-ME C_4_ grass, dicarboxylate transporter-2 (*ZmDCT2*, GRMZM2G086258) moves malate into the chloroplast of BS cells during C_4_ photosynthesis (Ding *et al*., 2015). *ZmDCT2* plays a critical role during C_4_ photosynthesis in *Z. mays*, and its absence severely impairs plant growth and development (Ding *et al*., 2015). The role of DCTs in C_4_ photosynthesis in other species, however, remains unknown.

*Z. mays* is the best characterized and functionally annotated C_4_ grass species. As such, it is a useful reference for identification of photosynthesis-related genes in poorly characterized C_4_ grasses and for resolving orthology (John *et al*., 2014; Ding *et al*., 2015; Huang *et al*., 2017). Microsynteny, the comparison of collinearity among related species, is a reliable approach to determine orthology and predict the function of a gene (Bennetzen and Freeling, 1997; Chen *et al*., 1997; Tikhonov *et al*., 1999; Bennetzen, 2000; Kumar *et al*., 2009; Jin *et al*., 2016). Davidson *et al.* (Davidson *et al*., 2012) showed that syntenic orthologs are likely to have conserved functions and expression patterns across lineages. Here we identify a new member of the *DCT* family, *DCT4*, which is not syntenic to the photosynthetic gene *ZmDCT2* and is not detected in the maize reference genome. We demonstrate that *Sorghum bicolor* DCT4 (SbDCT4) efficiently exchanges malate across membranes, consistent with a malate transport role in C_4_ photosynthesis. We characterize the diverse expression patterns of *DCT* genes in leaves of multiple 2 grass species. We also propose that sub-functionalization of *DCT*s in grasses of the PACMAD clade (Sanchez-Ken and Clark, 2010) occurred during independent recurrent evolution of C_4_ photosynthesis.

## RESULTS

### Identification of *DCT4* in *Sorghum bicolor*

To learn more about C_4_-related dicarboxylate transporters in species evolutionarily related to maize, we identified the syntenic ortholog of *ZmDCT2* in *S. bicolor*. Two genes, Sobic.007G226700 and Sobic.007G226800, are present at the predicted syntenic orthologous position on chromosome 7. We refer to them as *SbDCT2.1* and *SbDCT2.2*, respectively (Figure 1). *ZmDCT2* is abundantly expressed (Table 1), and its expression is enriched in BS cells of maize leaves (Figure 2) (Li *et al*., 2010; Tausta *et al*., 2014; Ding *et al*., 2015). In contrast, the expression profiles of *SbDCT2.1* and *SbDCT2.2* in *S. bicolor* leaves are low (Table 1). *SbDCT2.1* expression is slightly enriched in the M cells whereas *SbDCT2.2* is enriched in BS cells (Figure 2). We also analyzed the transcript levels of two other *S. bicolor* dicarboxylate transporters, *SbDCT1* (Sobic.002G233700) and *SbOMT1* (Sobic.008G112300). These genes are the orthologs of the *Z. mays* genes *ZmDCT1* (GRMZM2G040933) and *Zm-*oxoglutarate/malate transporter 1 (*ZmOMT1*; GRMZM2G383088), respectively. We found that *SbDCT1* expression, similar to that of *ZmDCT1*, is relatively low (Table 1), and only slightly differentially expressed in M cells relative to BS cells (Figure 2). The expression of both *ZmOMT1 and SbOMT1* is relatively high (Table 1), and both are slightly enriched in M cells (Figure 2).

**Table 1.**
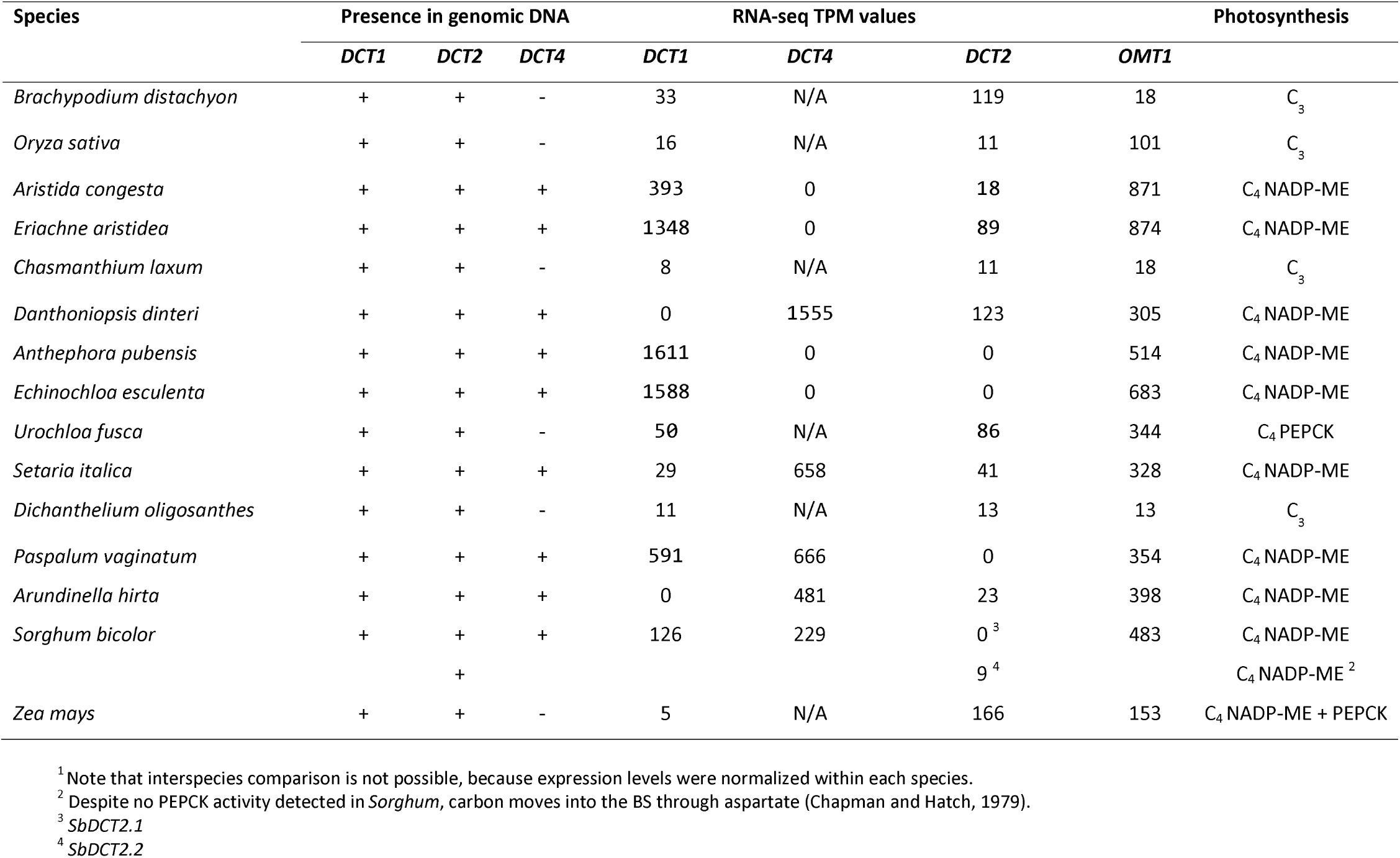
Genomic presence or absence and whole leaf TPM values for dicarboxylate transporter genes in grass leaves^1^.

**Figure 1.**
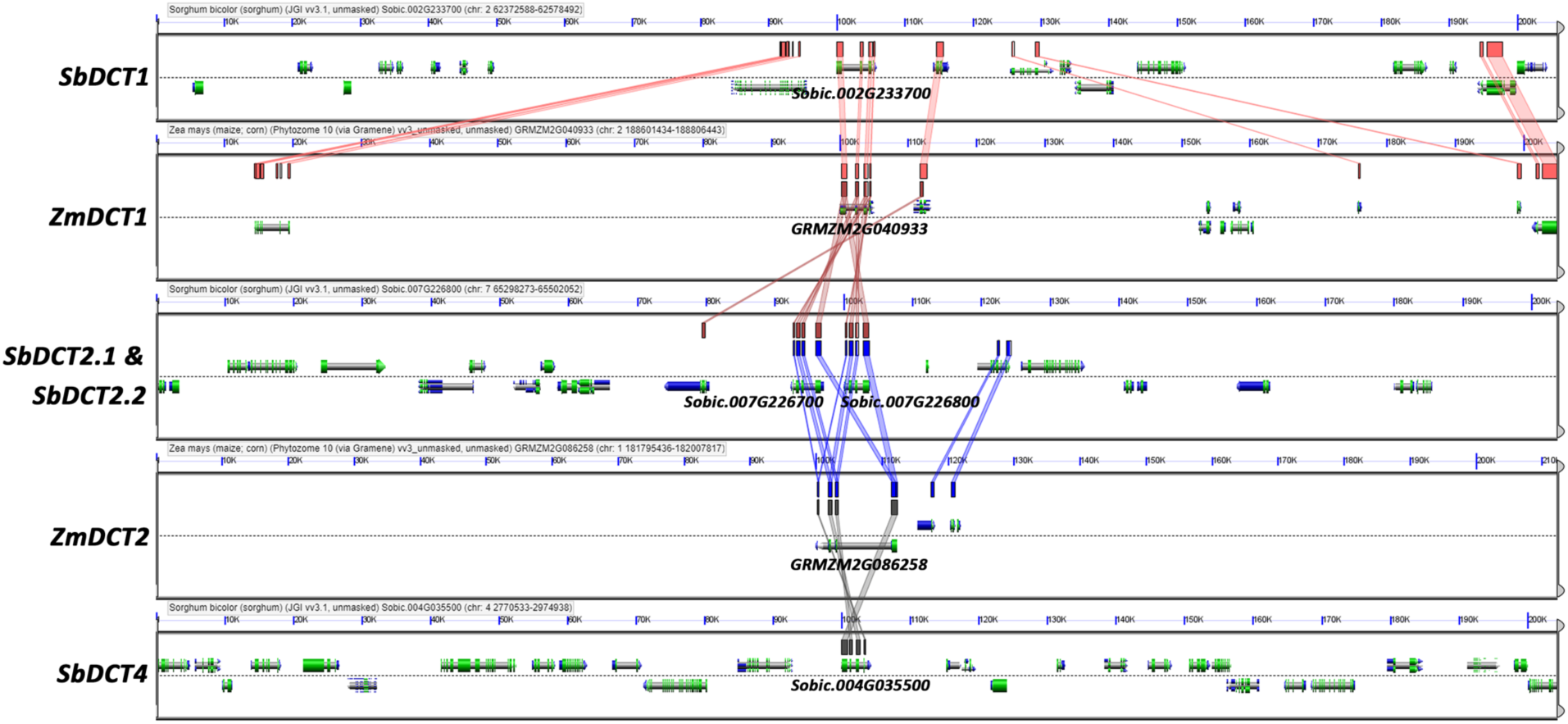
CoGe (https://genomevolution.org/coge/) genome viewer screenshots depicting the conservation and genomic contexts of *DCT* genes in *Z. mays* and *S. bicolor*. Colored lines between panels show conserved genes. *SbDCT4* shows high sequence conservation with other DCT genes, but is not a syntenic ortholog of *ZmDCT2*, as shown by the lack of conservation in neighboring genes.

**Figure 2.**
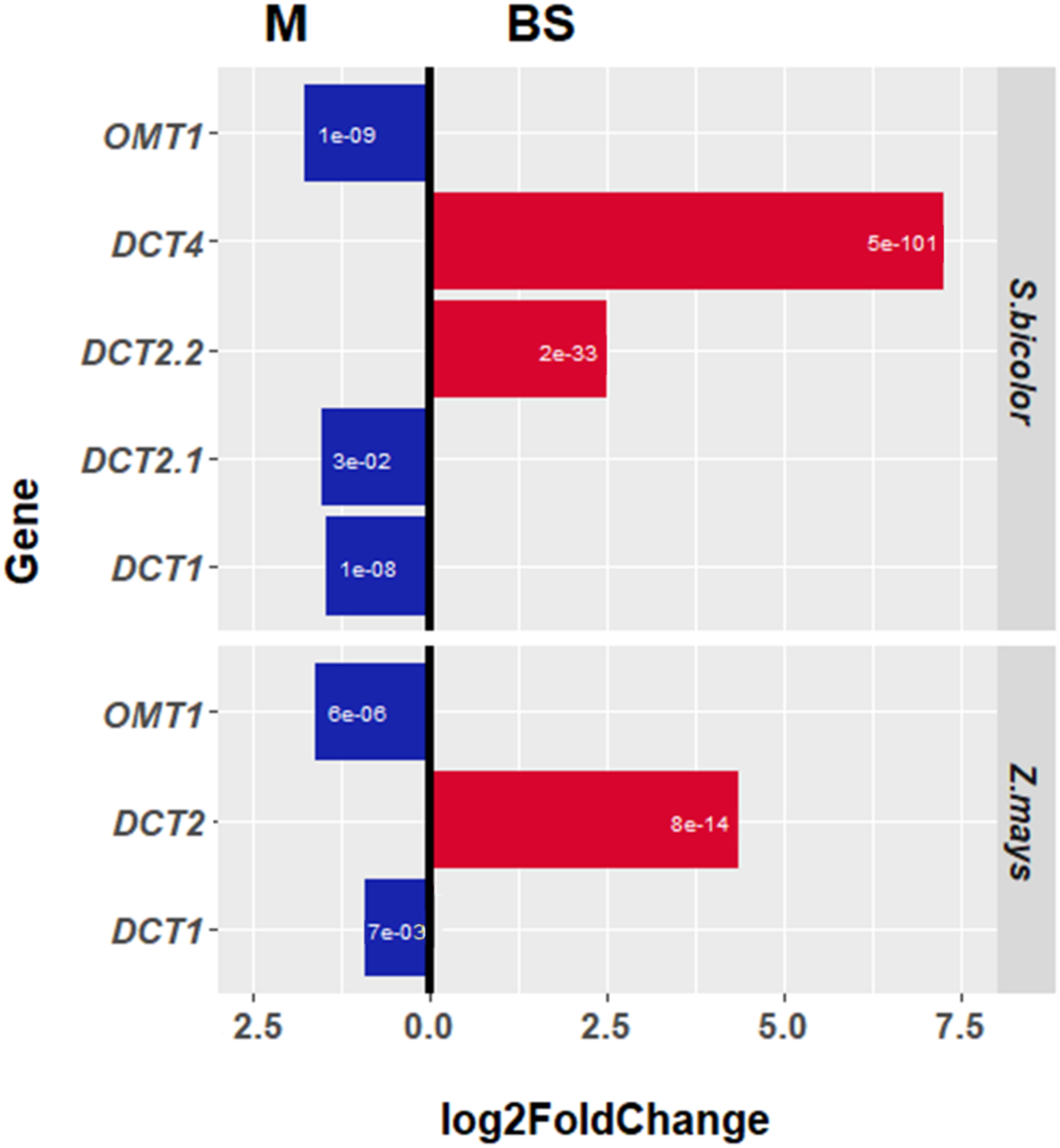
Differential expression of malate transporters in *Z. mays*, and *S. bicolor* leaves between the BS and M cells. The genome of *Z. mays* has one copy of *DCT2* and does not contain *DCT4*, and *ZmDCT2* is highly enriched in BS cells. *Sorghum bicolor* has two copies of *DCT2* (*DCT2.1*, and *DCT2.2*) in the syntenic genomic location that are the result of gene duplication. *S. bicolor* also expresses *DCT4*, which is highly enriched in BS cells. Both species express *OMT1* and *DCT1, which* are only slightly enriched in the M cells. Red bars represent enrichment in the BS cells. Blue bars represent enrichment in the M cells. The white numbers inside the bars represent the significance (p-value) of the log2(FoldChange).

In C_4_ species, the expression of many photosynthetic genes is enriched in either BS and M cells (Li et al., 2010; John et al., 2014; Tausta et al., 2014; Weissmann et al., 2015; Rao et al., 2016). In NADP-ME species, two transporters, one within the BS cells and another in M cells, move malate in and out of the chloroplast during C_4_ photosynthesis (Brautigam *et al*., 2008; Weissmann and Brutnell, 2012; John *et al*., 2014; Tausta *et al*., 2014; Wang *et al*., 2014a). However, in sorghum leaves, we found only one highly expressed dicarboxylate transporter, *SbOMT1*, that showed slightly enriched expression in M cells. Therefore we screened the sorghum genome for additional homologs of known maize *DCT*s. We identified the gene Sobic.004G035500 that showed homology to *ZmDCT1* and *ZmDCT2* but was not syntenic to either gene (Figure 1). *DCT3* is the name of the second transcript of *ZmDCT2* (Taniguchi *et al*., 2004), so we named this new gene *SbDCT4*. No syntenic ortholog of *SbDCT4* is present in the reference genomes of *Z. mays* or *Oryza sativa*. The absence of syntenic conservation between *S. bicolor* and *Z. mays* and the lack of direct orthologs in *Z. mays* or C_3_ species prevented identification of *DCT4* in a 3 previous bioinformatic screen for C_4_ photosynthesis genes (Huang *et al*., 2017). The expression of *SbDCT4* is moderately abundant (Table 1) and strongly enriched in the BS cells of *S. bicolor* leaves (Figure 2).

### SbDCT4 is an efficient malate transporter

To verify the ability of SbDCT4 to transport malate, we cloned coding sequences from the three sorghum *DCT* genes, *SbDCT1, SbDCT2*, and *SbDCT4*. We measured the malate transport activities of the recombinant proteins expressed in yeast. SbDCT4 was an efficient malate transporter (Table 2). The *K*_m_ of SbDCT4 was similar to that of SbDCT1, and the affinity for malate was highest in SbDCT2 among the three SbDCTs (Table 2), consistent with the relative malate transport activities reported for maize DCT1 and DCT2 (Taniguchi *et al*., 2004).

**Table 2.** *K*_m_ of malate for recombinant DCT proteins^1^ demonstrates the ability of SbDCT4 to transport malate efficiently.

| | $K_m$ (mM) | | |
| --- | --- | --- | --- |
|  | DCT1 | DCT2 | DCT4 |
| *S. bicolor* | 1.24 ± 0.14 | 0.71 ± 0.10 | 1.13 ± 0.10 |
| *Z. mays*<sup>2</sup> | 1.1 ± 0.1 | 0.85 ± 0.44 | N/A |

<sup>1</sup> The values are the means of three independent experiments ± SE.
<sup>2</sup> Kinetic values from a previous report (Taniguchi *et al.*, 2004).

### Phylogenetic distribution of *DCT* genes in grasses

To understand the relationship of *SbDCT4* to other grass *DCT* genes, we searched the genomes of the grass species *Setaria italica, Urochloa fusca, Brachypodium distachyon*, and *Dichanthelium oligosanthes*. In *S. italica*, an NADP-ME C_4_ species, we identified a dicarboxylate transporter, Seita.9G375100, that showed no syntenic orthologous relationship with dicarboxylate transporter genes in other available grass genomes. Phylogenetic analysis showed that this gene clustered with *SbDCT4* but not with *SbDCT1* and *SbDCT2* (Figure 3). We designated this gene *SiDCT4*. We did not detect orthologs, syntenic or otherwise, in *U. fusca*, a PEPCK C_4_ species, or in the two C_3_ species. To expand the search for *DCT4* in other grasses currently lacking genome assemblies, we examined leaf-derived transcript assemblies for *Aristida congesta, Eriachne aristidea, Chasmanthium laxum, Danthoniopsis dinteri, Anthephora pubensis, Echinochloa esculenta, Paspalum vaginatum*, and *Arundinella hirta* (Huang *et al*., in preparation). We then used the predicted coding sequences of the *DCT* genes from available genomes and from the *de novo* leaf transcriptome assemblies to generate a phylogenetic tree of the *DCT* family. The resulting phylogeny shows that *DCT4* transcripts form a distinct subclade from the *DCT1* clade (Figure 3). The absence of *DCT4* transcript expression does not rule out the existence of the gene in the genome. We also used polymerase chain reaction (PCR) to survey for *DCT* genes in the genomes of grass species for which whole-genome assemblies were not available. We designed conserved primers (non-degenerate or minimally degenerate) to small regions unique to each of the three *DCT* genes using PrimaClade (Gadberry *et al*., 2005). We detected *DCT1* and *DCT2* in the genomes of all species tested (Table 1, 4 Supplemental Figure 1). *DCT4*, however, was detected only in the genomes of NADP-ME C_4_ species of the PACMAD clade, excluding *Z. mays* (Table 1, Supplemental Figure 2).

**Figure 3.**
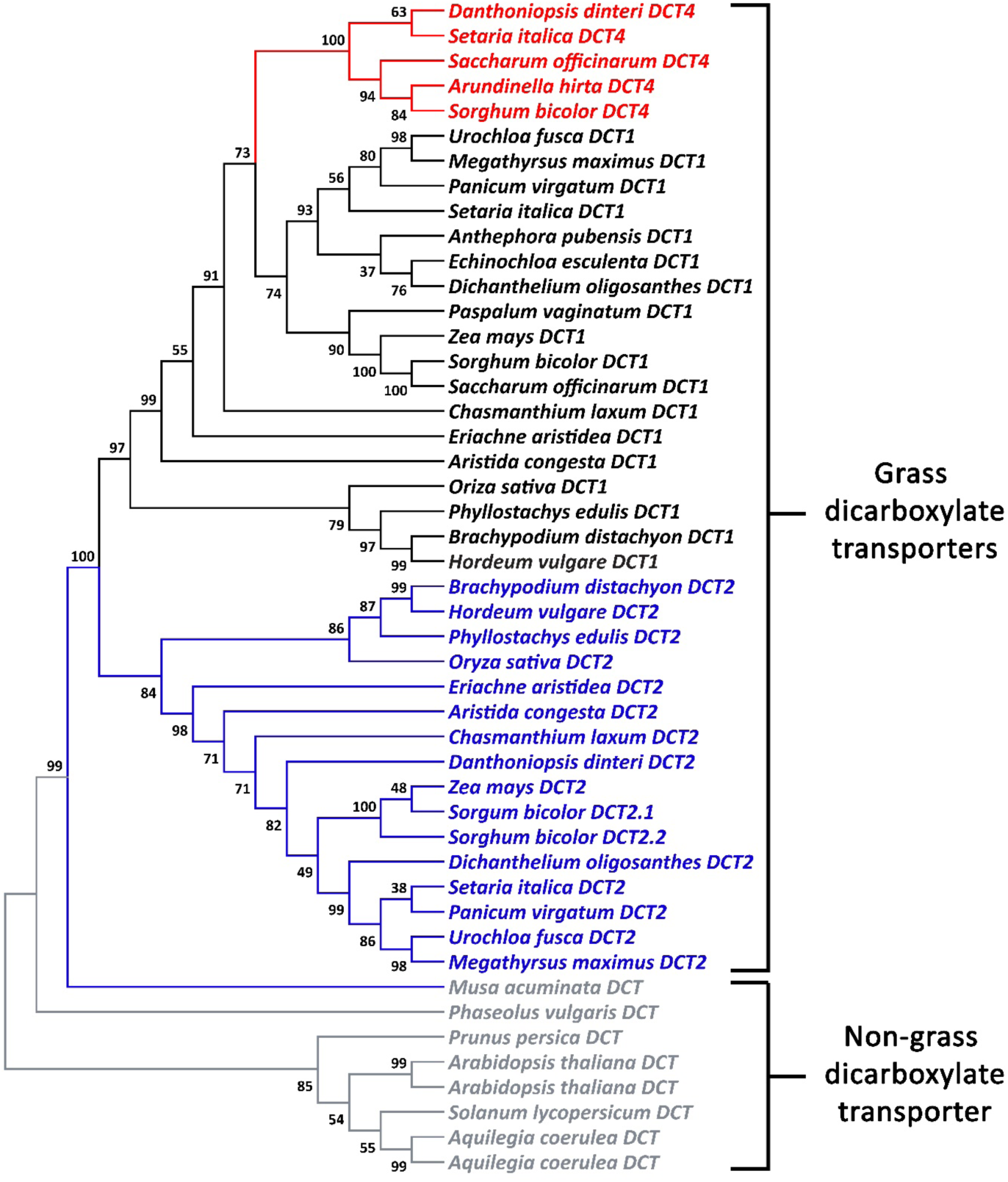
A phylogenetic tree of the *DCT* family in the grasses showing that *DCT4* is a subclade of *DCT1*. The *DCT1, DCT2*, and *DCT4* gene lineages are black, blue, and red, respectively. The length of the branches represents the evolutionary distance between ancestor to descendent nodes. The numbers represent the confidence level of the specific branch.

### Expression of malate transporter genes in NADP-ME C_4_ grasses

C_3_ species and *U. fusca*, a PEPCK C_4_ species, express both *DCT1* and *DCT2* at low levels in leaves (Table 1). C_4_ NADP-ME species of the PACMAD clade generally express one *DCT* gene in leaves at a high level and also express one or two other *DCT* genes at low levels (Table 1). We did not find an apparent lineage-specific pattern for the expression of the predominant *DCT* gene in the NADP-ME species we analyzed. This finding is consistent with random evolutionary processes underlying the sub-functionalization of members of the *DCT* family. Interestingly, *Z. mays* is the only species we examined in which *DCT2* is the predominantly expressed *DCT* gene (Table 1). We also examined the expression of the non-*DCT* malate transporter *OMT1* gene in the leaves of grasses (Table 1). Interestingly, we found that while *OMT1* expression was generally abundant, there was no consistent pattern of relative expression between the *DCT* and *OMT* genes within the NADP-ME C_4_ species (Table 1).

## DISCUSSION

### Evolution of the *DCT* gene family in grasses

We identified *DCT4* as a new member of the *DCT* gene family in the grasses (Figure 1). Our analysis suggests that *DCT4* is present in some C_4_ NADP-ME PACMAD grasses. *DCT1* and *DCT2* appear to have originated from a duplication of a single *DCT* gene after the monocot-eudicot split (Taniguchi *et al*., 2004) and *DCT4* arose from a duplication of *DCT1* at the root of the PACMAD grasses (Figure 3). The expression of *DCT* genes in the grasses that we analyzed exhibited no clear lineage-specific patterns (Table 1). Therefore, we propose that different members of the *DCT* family subfunctionalized for photosynthetic malate transport in the BS cells of C_4_ grasses of the PACMAD clade.

This work also confirms the importance of including syntenic and expression data in assigning orthology across species, and of developing multiple models for C_4_ photosynthesis in the grasses. *SiDCT4* was previously mis-annotated as the ortholog of *DCT2* (John *et al*., 2014), likely because of the lower expression level of *SiDCT2* (Seita.9G375100) in leaf tissue. The use of different malate transporters, for example, DCT4 in *S. bicolor* and *S. italica*, or DCT2 in *Z. mays*, suggests that multiple evolutionary paths resulted in the development of an active C_4_ NADP-ME photosynthetic cycle. It is interesting to note 5 common origins of C_4_ photosynthesis are often defined based on the predominant decarboxylase utilized, thus maize and sorghum are considered to have evolved from a common C_4_ ancestor. This analysis suggests that rather than being static, biochemical adaptations continued after the divergence of maize and sorghum lineages. Thus, optimizations of C_4_ activities may be continuous as breeding pressures or climate change alters ecological niches of individual species.

### Various C_4_ subtype combinations have different transport requirements

The variation of expression levels among the different malate transporters within each NADP-ME species (Table 1) suggests different transport requirements during C_4_ photosynthesis. This supposition is in agreement with the view that the three subtypes of C_4_ photosynthesis are mixed rather than exclusive (Hatch, 1971; Chapman and Hatch, 1979; Furbank, 2011; Pick *et al*., 2011; Wang *et al*., 2014b). For example, *Z. mays* utilizes both the NADP-ME (75%) and PEPCK (25%) pathways to fix carbon (Chapman and Hatch, 1979; Wingler et al., 1999; Weissmann et al., 2015), and has similar expression levels of *DCT2* and *OMT1* and low expression of *DCT1. S. bicolor* moves carbon through both malate and aspartate, although no PEPCK activity was detected in its leaves (Chapman and Hatch, 1979). *S. bicolor* has similar expression levels for *DCT4* and *DCT1* and high expression of *OMT1* (Table 1). Other grass species may have dicarboxylate transporter expression ratios that correspond to their unique combination of C_4_ subsystems. For example, *OMT1* is highly expressed in *U. fusca*, ∼3-7 fold higher than *DCT2* or *DCT1*, respectively. OMT1 transports dicarboxylates, excluding those containing an amino group (Taniguchi *et al*., 2002; Taniguchi *et al*., 2004). Thus, in PEPCK C_4_ plants, OMT1 may move oxaloacetate into the mesophyll chloroplast, and 2-oxoglutarate out, to support the high production of aspartate needed to maintain the photosynthetic cycle (Rathnam and Edwards, 1977). Interestingly, both *ZmOMT1* and *SbOMT1* are only slightly differentially expressed in the M cells (Figure 2). As the loss of *DCT2* in *Z. mays* prevents movement of malate into the BS chloroplast (Weissmann et al., 2015), *OMT1* cannot be moving malate into the BS chloroplast alongside *DCT2*. But *OMT1* may also have a role in organic acid metabolism in both cell types, such as shuttling reducing equivalents in organelles other than the chloroplast (Pleite *et al*., 2005).

## CONCLUSIONS

Our results show that the newly identified member of the *DCT* gene family, *SbDCT4*, is an efficient malate transporter. Based on the expression patterns of malate transporters among the grasses, we suggest that different members of the *DCT* family may have evolved multiple roles in C_4_ photosynthesis. 6 Further studies will be needed to verify the subcellular localization of these proteins and to define their specific metabolic functions. Characterizing the various combinations of C_4_ photosynthetic subsystems in grasses will facilitate the exploitation of *DCT* genes, through breeding or engineering, to improve the performance of crop plants and increase yield.

## MATERIALS AND METHODS

### Identification of *DCT4* genes in *Sorghum* and other grasses

We used QUOTA-ALIGN (Tang *et al*., 2011) to identify syntenic orthologous regions in grass species with sequenced genomes, following the protocol described in (Zhang et al., 2017). To find homologous genes at non-syntenic locations, we used two complementary approaches. For species with sequenced genomes, we used LASTZ (Harris, 2007) to align the coding sequence of the primary transcript annotated in Phytozome (https://phytozome.jgi.doe.gov) to the genome assembly. For species without assembled genomes, we used LASTZ to align the coding sequence of the primary transcript from Phytozome to transcript assemblies generated by Trinity (Grabherr *et al*., 2011).

### Measurements of malate transport

We cloned each of the three *SbDCT* cDNAs between the promoter and terminator of yeast *GAL2* in the pTV3e vector (Nishizawa *et al*., 1995). We transformed the plasmids into yeast LBY416 cells and selected transformants on tryptophan-deficient agar plates. We prepared a crude membrane fraction from the selected yeast transformants. We used a freeze-thaw technique to reconstitute liposomes for the measurement of the uptake of [^14^C]malate (Taniguchi *et al*., 2002).

### Phylogenetic analysis of *DCT* homologs

*DCT* coding sequences for *Z. mays, S. bicolor, S. italica, B. distachyon, O. sativa, D. oligosanthes*, and *U. fusca* were from Phytozome (https://phytozome.jgi.doe.gov). We used BLASTN (Altschul *et al*., 1990) to search *de novo* assembled leaf transcriptomes (Huang, manuscript in preparation) from the C_4_ grass species *A. congesta, E. aristidea, C. laxum, D. dinteri, A. pubensis, E. esculenta, P. vaginatum*, and *A. hirta* with the *DCT* sequences from maize, *Setaria*, and *Sorghum* as queries. We used ProGraphMSA to generate a codon-based sequence alignment (Szalkowski, 2012). We used MEGA6 (Tamura *et al*., 2013), with default parameters and the branch support values based on 1,000 bootstraps, to generate the phylogenetic reconstruction with the maximum likelihood method and based on the nucleotides in the third position of codons (Simmons *et al*., 2006).

### Analysis of gene expression for decarboxylase transporters in grasses

For species with published leaf transcriptome profiles (Ouyang *et al*., 2007; Li *et al*., 2010; Zhang *et al*., 2012; Schnable, 2014; Wang *et al*., 2014a; Studer *et al*., 2016), gene expression levels were calculated and normalized, for each species, as Transcripts Per Million (TPM). For the other species, the normalized TPM values were based on *de novo* transcriptome assemblies (Huang *et al*., manuscript in preparation). The values in Table 1 only allow for intraspecies comparisons among the decarboxylase transporters.

### Identification of *DCT4* in species without sequenced genomes

We aligned the coding sequences from each of the *DCT* genes from *Z. mays, S. bicolor, S. italica, U. fusca, B. distachyon, O. sativa, D. oligosanthes, A. congesta, E. aristidea, C. laxum, D. dinteri, A. pubensis, E. esculenta, P. vaginatum*, and *A. hirta* using PAL2NAL (Suyama *et al*., 2006). The resulting multiple sequence alignment enabled the design of non-degenerate or minimally degenerate PCR primers (Table 3) using PrimaClade (Gadberry *et al*., 2005).

**Table 3.**
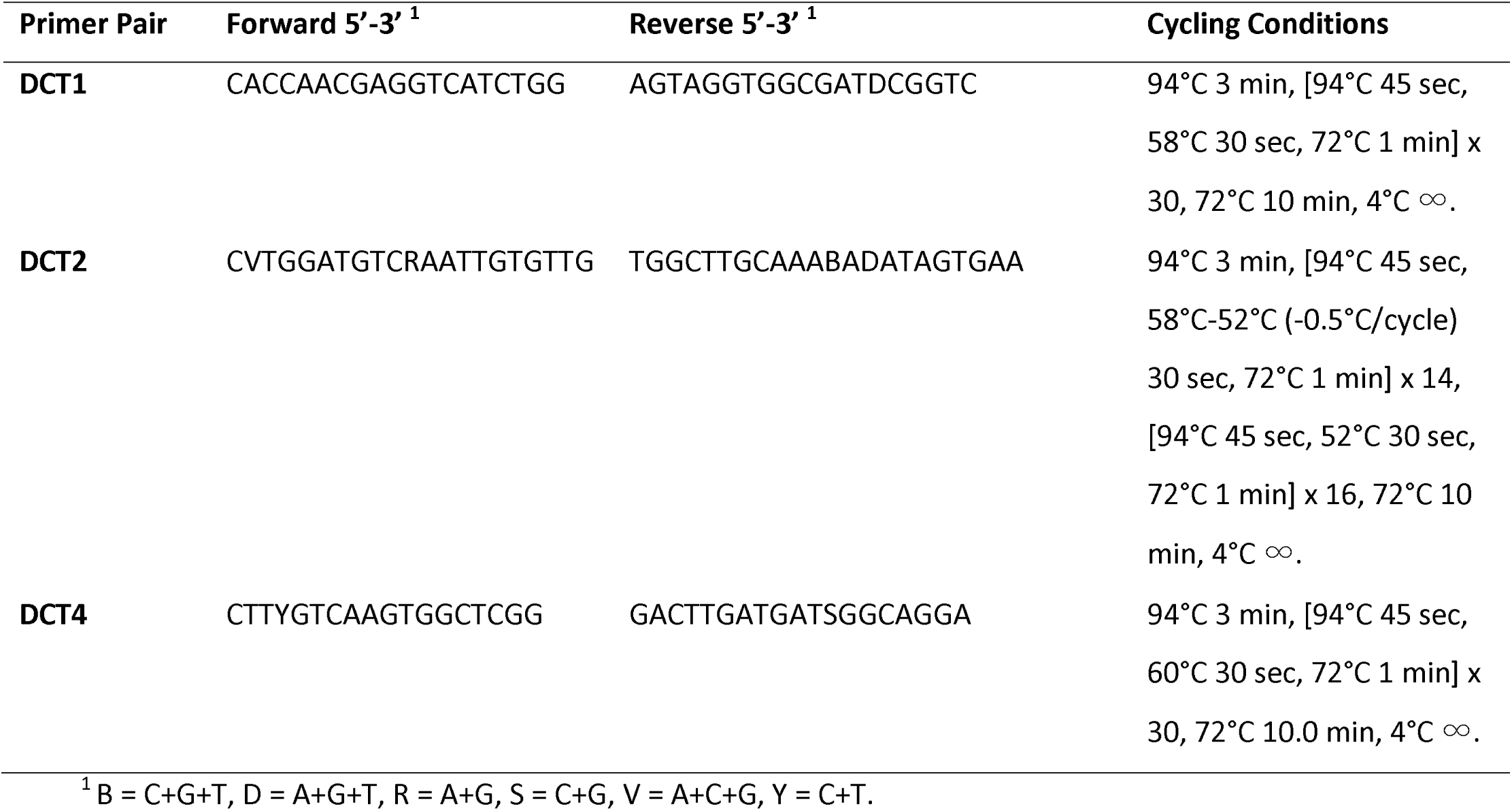
Primers and PCR conditions for the amplification of grass *DCT* genes.

Jacob D. Washburn and J. Chris Pires (University of Missouri, Columbia) kindly provided genomic DNA from *A. congesta, E. aristidea, D. dinteri, A. pubensis, E. esculenta*, and *A. hirta* (Washburn *et al*., 2015). We used a CTAB-based method to extract genomic DNA from *C. laxum, P. vaginatum, Z. mays, S. bicolor, S. italica*, and *B. distachyon* (Weissmann *et al.*, 2015). *Z. mays* and *B. distachyon* were the negative controls for *DCT4* and the positive controls for *DCT1* and *DCT2. S. bicolor* and *S. italica* were the positive controls for *DCT1, DCT2*, and *DCT4.*

We conducted amplification of *DCT* genes by PCR using a 25-μl reaction mix and an ABI 2720 Thermal cycler. The reaction mixture included 2.5 μl of 10X Buffer, 2.5 μl of 10 μM solutions of forward and reverse primers, 2 μl of 2.5 mM dNTP stock, 14 μl of nuclease-free water, 0.5 μl of Choice Taq enzyme, and 1 μl of 100 ng/μl DNA. We performed PCR reactions as described in Table 3 with 5 μl of loading dye added to each reaction. Aliquots of 13 μl were loaded on 3% agarose gels (Invitrogen UltraPure Agarose 1000, 1X TAE buffer, Invitrogen SYBR Safe Gel Stain) and electrophoresed for 30 minutes at 100 volts. We based size estimates on 100bp and 50 bp DNA markers (GoldBio). 8

## FIGURE LEGENDS

**Supplemental Figure 1.**
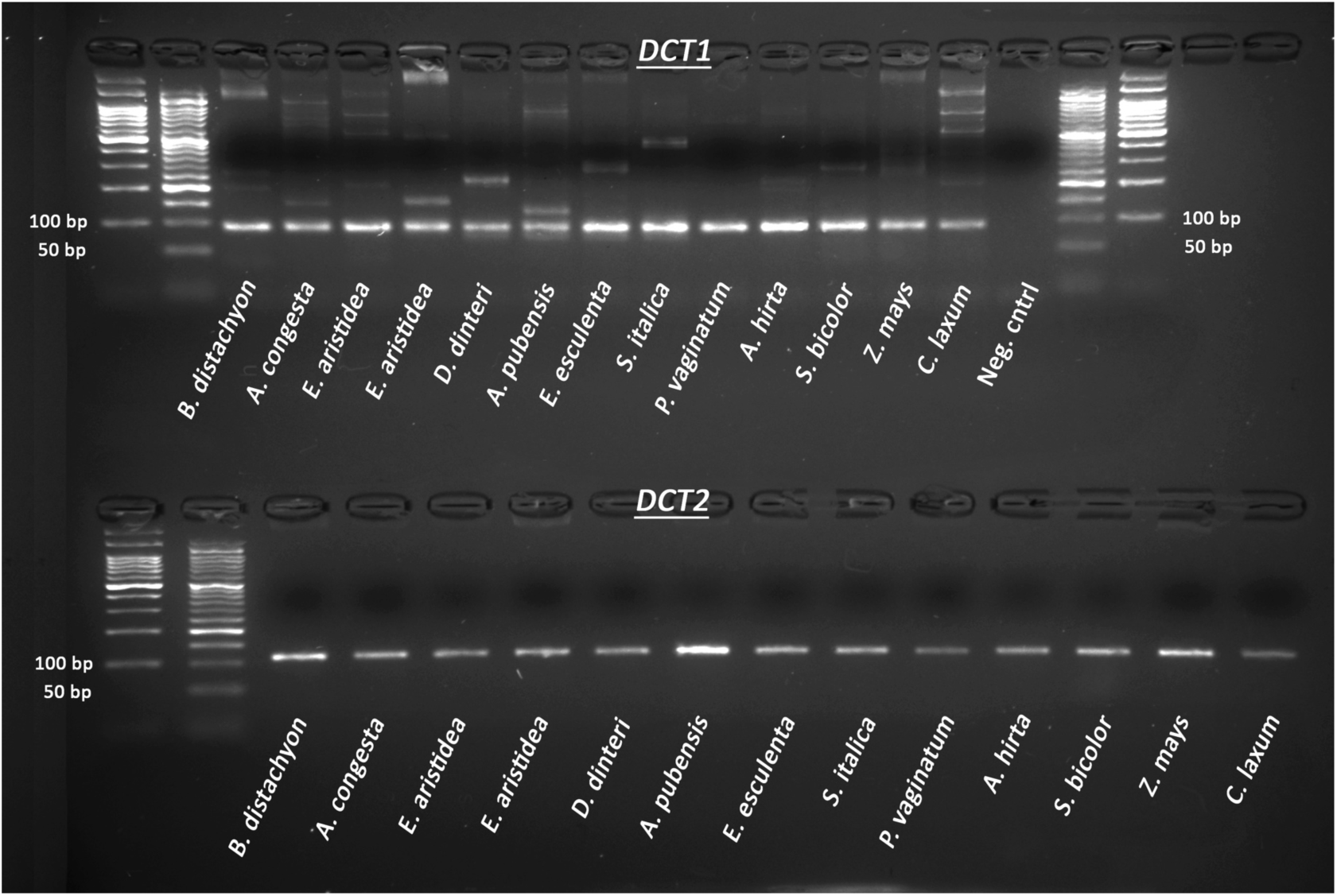
Gel image showing that *DCT1* and *DCT2* genes are present in all grass species tested.

**Supplemental Figure 2.**
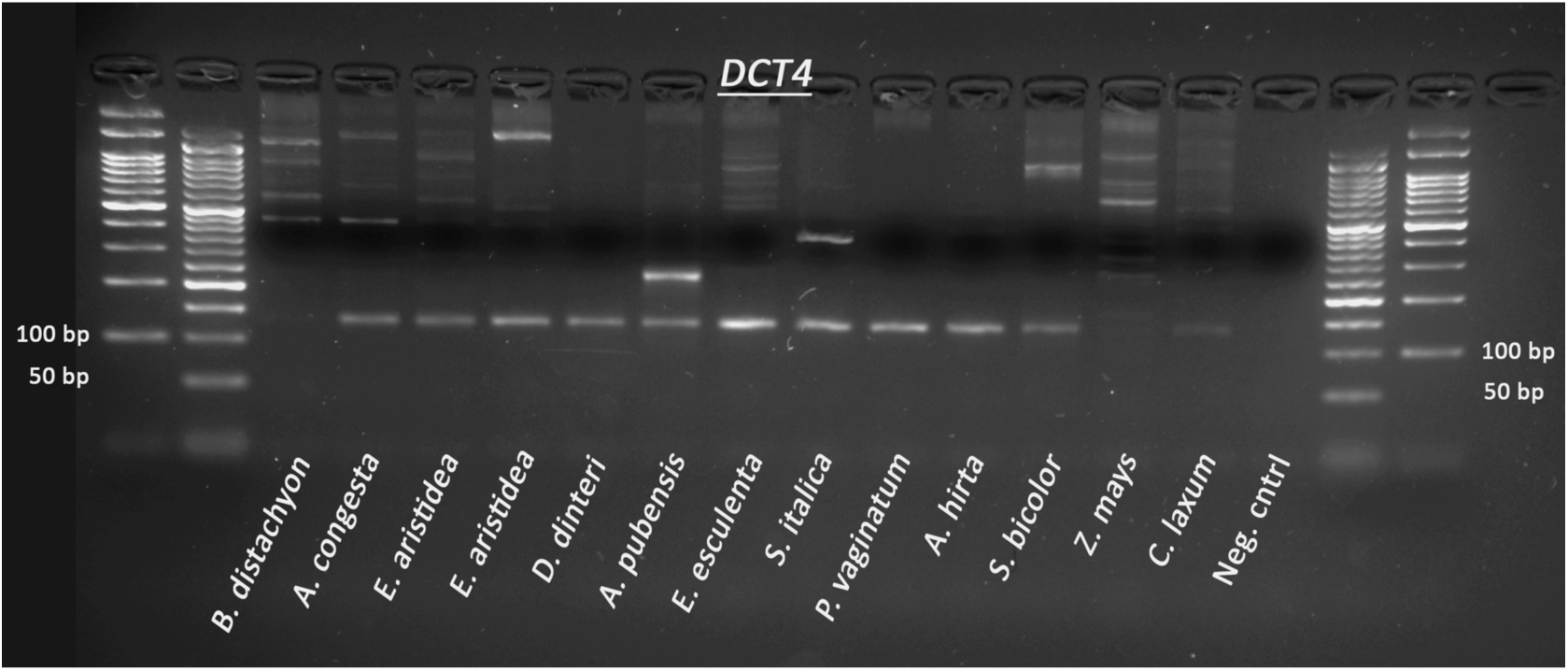
Gel image showing the presence or absence of *DCT4* genes from species lacking genome assemblies. Negative controls were *Z. mays* and *B. distachyon*, and positive controls were *S. bicolor* and *S. italica*.

